# The Relationship of *pqs* Gene Expression to Acylhomoserine Lactone Signaling in *Pseudomonas aeruginosa*

**DOI:** 10.1101/2024.03.22.586172

**Authors:** Martin P. Soto-Aceves, Nicole E. Smalley, Amy L. Schaefer, E. Peter Greenberg

## Abstract

The opportunistic pathogen *Pseudomonas aeruginosa* has complex quorum sensing (QS) circuitry, which involves two acylhomoserine lactone (AHL) systems, the LasI AHL synthase and LasR AHL-dependent transcriptional activator system and the RhlI AHL synthase-RhlR AHL-responsive transcriptional activator. There is also a quinoline signaling system (the *Pseudomonas* quinolone signal, PQS, system). Although there is a core set of genes regulated by the AHL circuits, there is substantial strain-to-strain variation in the non-core QS regulated genes. Reductive evolution of the QS regulon, and variation in specific genes activated by QS, occurs in laboratory evolution experiments with the model strain PAO1. We used a transcriptomics approach to test the hypothesis that reductive evolution in the PAO1 QS regulon can in large part be explained by a simple null mutation in *pqsR*, the gene encoding the transcriptional activator of the *pqs* operon. We found that PqsR had very little influence on the AHL QS regulon. This was a surprising finding because the last gene in the PqsR-dependent *pqs* operon, *pqsE*, codes for a protein, which physically interacts with RhlR and this interaction is required for RhlR-dependent activation of some genes. We used comparative transcriptomics to examine the influence of a *pqsE* mutation on the QS regulon and identified only three transcripts, which were strictly dependent on PqsE. By using reporter constructs we showed that the PqsE influence on other genes was dependent on experimental conditions and we have gained some insight about those conditions. This work adds to our understanding of the plasticity of the *P. aeruginosa* QS regulon and to the role PqsE plays in RhlR-dependent gene activation.

## INTRODUCTION

The opportunistic human pathogen *Pseudomonas aeruginosa* is a model for studies of quorum sensing (QS) (Schuster *et al*., 2013). There are two acylhomoserine lactone (AHL) QS circuits, which together regulate expression of dozens to hundreds of genes depending on the strain under investigation. These are the LasR-LasI circuit and the RhlR-RhlI circuit. LasR is a transcriptional activator that responds to the QS signal synthesized by LasI, N-(3-oxododecanoyl)-L-homoserine lactone (3OC12-HSL). RhlR is a transcriptional activator that responds to the QS signal synthesized by RhlI, *N*-butanoyl-L-homoserine lactone (C4-HSL). In some strains, including the one we use in this study, PAO1, *rhlR* and *rhlI* expression require activation by LasR. The two systems regulate overlapping sets of genes, many of which code for virulence factors (Schuster *et al*., 2003, Wagner *et al*., 2003). There is an additional non-AHL QS system, the *Pseudomonas* quinolone signal (PQS) system, which interacts with the AHL QS systems in complicated ways [for reviews see (Trottier *et al*., 2024, Heeb *et al*., 2011)].

There is substantial strain-to-strain variation in the size of the QS regulon, but a core set of 43 genes, which are activated by QS in most strains examined has been identified (Chugani *et al*., 2012). Direct targets of LasR in strain PAO1 have been identified by ChiPseq (Gilbert *et al*., 2009), and recently, a set of direct RhlR targets in strain PA14 was identified by RNAseq (Keegan *et al*., 2023). In strain PAO1 there are hundreds of QS-activated genes (Schuster *et al*., 2003, Wagner *et al*., 2003). The census of QS-activated genes in PAO1 will vary depending on the experimental conditions employed. A simple approach to capturing the bulk of QS-activated genes in RNAseq experiments is a comparative transcriptomics approach where RNA is isolated from cells grown at 37 C aerobically in standard culture media (buffered Lennox broth) to late logarithmic phase. Transcriptomes of QS mutants can be compared with their wildtype (WT) parent or transcriptomes of WT grown in broth supplemented with a signal-inactivating lactonase can be compared to untreated cells (Schuster *et al*., 2003, Chugani *et al*., 2012, Smalley *et al*., 2022). These approaches do not capture the entire repertoire of QS-activated genes, some of which are silenced in late logarithmic phase and some of which require other conditions and are activated only later in stationary phase. Nevertheless, this experimental design captures a large majority of QS-activated genes, about 150-190 genes in PAO1 show QS activation over 3-fold (range from 3-fold to over 60-fold) in these sorts of analyses (Schuster *et al*., 2003, Chugani *et al*., 2012, Smalley *et al*., 2022).

The production of PQS is induced by PQS (or the PQS precursor 2-heptyl-4-quinolone) and the transcriptional activator PqsR (Gallagher *et al*., 2002) [also called MvfR (Cao *et al*., 2001)]. The *pqsR* gene is among the genes activated by LasR (Schuster *et al*., 2003). PqsR activates transcription of its linked *pqsA-E* operon and in stationary phase *pqsR* mutants express many AHL-dependent genes at lower levels than WT (Xiao *et al*., 2006, Cao *et al*., 2001). The *pqsA-D* genes are required for PQS synthesis and the last gene in the operon, *pqsE* is not required for PQS synthesis (Gallagher *et al*., 2002, Déziel *et al*., 2004). Rather PqsE has been shown to serve as a chaperone for RhlR (Groleau *et al*., 2020, Borgert *et al*., 2022, García-Reyes *et al*., 2021, Taylor *et al*., 2021). We note that most studies of the relationship between PqsE and RhlR have utilized strain PA14 and not PAO1. Some RhlR-dependent genes are silent in PA14 PqsE mutants and activation of others is partially suppressed in PqsE mutants (Keegan *et al*., 2023).

In a recent report we described a large reduction in the PAO1 QS regulon when cells were grown for about 1000 generations in media containing casein and adenosine (Smalley *et al*., 2022). Growth on either of these carbon sources requires QS activation of genes for their metabolism, and although LasR-LasI and RhlR-RhlI remain functional in this growth regime, many QS-activated genes not involved in adenosine or casein metabolism are silenced, or at least no longer QS-activated, in evolved populations. Isolates with *pqsR* mutations or with mutations that reduced and delayed *pqsR* expression were common in the evolved populations (Smalley *et al*., 2022). This finding leads to a hypothesis that a simple mutation in *pqsR* or affecting *pqsR* expression could be responsible for the bulk of the reduction in the size of the QS operon in these experiments, either by loss of PQS or by loss of PqsE. Here we show that the large reduction in QS-regulon size described previously (Smalley *et al*., 2022) does not result from a single gene mutation in either *pqsR* or *pqsE*. In addition, our transcriptomics and transcription reporter experiments show only three transcriptional units for which RhlR activity is essentially dependent on PqsE under the conditions of our transcriptomics experiments. By using transcriptional reporters we also begin to define the conditions where RhlR activity is stimulated by PqsE.

## RESULTS

### Transcriptomics show the QS regulon of WT PAO1 and a PqsR mutant are similar

Long-term growth of WT *P. aeruginosa* on casein and adenosine leads to a reduction of the number of genes activated by QS (30% reduction in one evolved population and 60% in another). Genomic sequencing of isolates from these populations showed that among several mutations, those impairing or eliminating *pqsR* expression were common (Smalley *et al*., 2022). This finding leads to the suggestion that loss of PqsR, or the last gene in the PqsR-activated *pqs* operon, *pqsE*, could provide a simple explanation for the large reduction in the number of genes activated by QS in the evolved populations. To test the hypothesis that a *pqsR* mutation could provide an explanation for at least much of the reduction in the QS regulon, we compared the QS regulon of WT to that of a *pqsR* deletion mutant by precisely the method used to identify the QS-activated genes in the reductive evolution report (Smalley *et al*., 2022). As in the previous report any gene, which showed a 2.8-fold (1.5-fold log_2_) or greater transcript level in cells grown without an inactivating lactonase as compared to cells grown with the lactonase were called as QS-activated (Table 1). As in the previous report cells were harvested for RNAseq analysis in late logarithmic growth phase [as determined using a Genesys spectrophometer (ThermoFisher, Waltham, MA) at an optical density at 600 nm (OD_600_) reading of 2]. The one difference between our experiment and the one reported previously (Smalley *et al*., 2022) is that cells used to inoculate buffered LB for the previous RNAseq experiments were from PAO1, which had been passaged daily for five days in casein plus adenosine broth. In our experiments, cells were grown only in buffered LB. Of the 129 genes we called QS-activated, 101 were identified as QS-activated in cells grown from 5-day casein plus adenosine broth cultures (Table 1). We believe the results to be in reasonably good concordance with the previous report. The differences might result from the culture histories of the inocula or, in some cases, the inherent error in the RNAseq procedure (particularly for genes just above the threshold of 2.8-fold change in cells grown without lactonase as compared to those grown with lactonase).

**Table 1.** Comparison of genes activated by QS in WT, Δ*pqsR*, and reduced regulon populations.

| Locus | Gene name | Gene activation (lactonase untreated vs. treated fold-change) |  |  |  |
| --- | --- | --- | --- | --- | --- |
| | | WT | $\Delta pqsR$ mutant | 5-day adenosine-casein pop<sup>a</sup> | 160 casein-adenosine pop<sup>a</sup> |
| PA0026<sup>b</sup> | *plcB* | 4.5 | 4.4 | 4.9, 3.4 | -, 3.6 |
| PA0027 |  | 4.0 | 5.4 | 5.5, 4.0 | -, 5.0 |
| PA0028 |  | 3.2 | 6.0 | 4.4, 4.4 | -, 3.9 |
| PA0052 |  | 6.7 | 6.6 | 10.4, 5.6 | 4.3, 6.4 |
| PA0105 | *coxB* | 3.0 | 5.3 | -, - | -, - |
| PA0106 | *coxA* | 3.9 | 6.7 | -, - | -, - |
| PA0107 |  | 3.1 | 6.1 | -, - | -, - |
| PA0108 | *colI* | 3.5 | 6.6 | -, - | -, - |
| PA0122 | *rahU* | 3.7 | 3.6 | 17.2, 11.0 | 3.1, 5.5 |
| PA0143 | *nuh* | 7.0 | 5.7 | 3.6, 2.9 | -, - |
| PA0144 |  | 7.1 | 6.7 | 5.1, 6.9 | -, - |
| PA0572 | *impA* | 15.4 | 22.5 | 5.6, 3.2 | -, 3.1 |
| PA0852 | *cbpD* | 18.3 | 14.9 | 17.3, 12.8 | 3.4, 7.0 |
| PA0855 |  | 2.9 | 2.9 | 4.4, 3.1 | -, 3.3 |
| PA0996 | *pqsA* | 6.7 | 1.2<sup>c</sup> | 41.2, 22.6 | -, - |
| PA0997 | *pqsB* | 9.1 | -1.1 | 35.4, 14.3 | -, - |
| PA0998 | *pqsC* | 5.0 | 1.2 | 15.5, 5.2 | -, - |
| PA0999 | *pqsD* | 2.9 | 1.1 | 8.1, - | -, - |
| PA1002 | *phnB* | 5.8 | 6.5 | 4.6, 5.1 | -, 3.3 |
| PA1003 | *pqsR* | 7.9 | 1.2<sup>d</sup> | 3.7, 3.9 | -, 3.0 |
| PA1168 |  | 85.2 | 179.7 | 33.3, 46.1 | 66.3, 9.3 |
| PA1169 |  | 14.4 | 18.2 | -, - | 5.1, - |
| PA1216 |  | 2.9 | 3.5 | 9.0, 4.6 | 3.5, 5.1 |
| PA1218 |  | 2.8 | 3.0 | -, - | -, - |
| **PA1221<sup>e</sup>** |  | 3.4 | 3.6 | 8.7, 6.4 | 6.6, 10.6 |
| PA1245 | *aprX* | 11.9 | 9.1 | 18.4, 15.7 | 4.4, 10.1 |
| PA1246 | *aprD* | 13.8 | 12.4 | 21.9, 16.5 | 4.3, 8.0 |
| PA1247 | *aprE* | 17.1 | 10.9 | 18.1, 14.9 | 4.3, 8.9 |
| PA1248 | *aprF* | 14.2 | 10.1 | 17.1, 14.1 | 3.9, 8.9 |
| PA1249 | *aprA* | 30.4 | 23.4 | 25.6, 19.0 | 6.3, 10.4 |
| PA1250 | *aprl* | 16.8 | 19.2 | 17.3, 21.4 | 3.2, 22.8 |
| PA1251 |  | 7.1 | 5.7 | 11.6, 11.8 | -, 12.6 |
| PA1431 | *rsaL* | 18.7 | 27.2 | 25.4, 20.5 | 4.1, 21.7 |
| PA1432 | *lasI* | 16.0 | 24.1 | 7.7, 8.7 | 3.2, 36.6 |
| PA1433 |  | 4.0 | 3.4 | -, - | -, 7.9 |
| PA1435 |  | 3.2 | 4.6 | -, - | -, - |
| PA1656 | *hsiA2* | 3.7 | 3.4 | 8.9, 8.5 | -, 8.4 |
| PA1657 | *hsiB2* | 3.7 | 4.7 | 9.2, 7.8 | -, 3.5 |
| PA1658 | *hsiC2* | 4.0 | 4.2 | 7.1, 5.3 | -, - |
| PA1659 | *hsiF2* | 3.2 | 3.3 | 10.2, 6.3 | -, 3.4 |
| PA1660 | *hsiG2* | 3.2 | 3.5 | 5.7, 4.2 | -, - |
| PA1664 | *orfX* | 3.8 | 3.8 | 6.0, 6.1 | -, - |
| PA1665 | *fha2* | 3.7 | 3.3 | 4.9, 6.1 | -, - |
| PA1784 |  | 3.7 | 4.4 | 9.1, 5.4 | -, 3.5 |
| **PA1869** | *acp1* | 6.5 | 7.3 | 38.5, 43.1 | 9.5, 13.1 |
| PA1871 | *lasA* | 8.9 | 12.2 | 7.2, 4.5 | -, 3.9 |
| PA1892 |  | 5.4 | 4.8 | 3.6, 4.1 | -, - |
| PA1893 |  | 7.7 | 6.9 | 3.0, - | -, - |
| PA1894 |  | 29.8 | 21.5 | 9.8, 10.8 | 3.0, - |
| PA1895 |  | 26.5 | 18.4 | -, - | -, - |
| PA1896 |  | 17.8 | 18.0 | 3.1, 2.8 | -, - |
| PA1897 |  | 36.3 | 34.8 | 5.2, 5.4 | -, - |
| **PA1899<sup>f</sup>** | *phzA* | 6.7 | 5.1 | -, - | -, - |
| **PA1900<sup>f</sup>** | *phzB* | 3.1 | 3.6 | -, - | -, - |
| PA2076 |  | 5.3 | 3.9 | 4.7, 4.5 | -, 3.8 |
| **PA2193** | *hcnA* | 8.7 | 10.8 | 29.0, 31.4 | 11.2, 23.3 |
| **PA2194** | *hcnB* | 8.0 | 9.4 | 11.8, 14.9 | 4.3, 7.1 |
| **PA2195** | *hcnC* | 7.7 | 6.4 | 8.6, 11.8 | 2.9, 3.9 |
| PA2300 | *chiC* | 4.0 | 3.3 | 4.4, 3.6 | -, - |
| PA2302 | *ambE* | 16.1 | 32.6 | 18.2, 19.7 | -, 6.9 |
| PA2303 | *ambD* | 38.2 | 42.4 | 23.4, 31.5 | 4.5, 11.7 |
| PA2304 | *ambC* | 21.8 | 41.6 | 16.4, 16.1 | -, 7.9 |
| PA2305 | *ambB* | 13.3 | 28.7 | 12.9, 16.1 | -, 6.5 |
| PA2366 | *hsiC3* | 2.9 | 5.2 | 3.1, - | -, - |
| PA2367 | *hcp3* | 2.9 | 5.4 | 3.6, - | -, - |
| PA2375 |  | 3.4 | 3.2 | -, - | -, - |
| PA2384 |  | 3.4 | 5.5 | -, - | -, - |
| PA2414 |  | 3.5 | 3.4 | -, - | -, - |
| PA2415 |  | 3.1 | 4.1 | -, - | -, - |
| PA2423 |  | 4.1 | 4.6 | 6.1, 4.7 | -, 3.5 |
| PA2476 | *dsbG* | 3.0 | 3.3 | -, - | -, - |
| PA2507 | *catA* | 3.3 | 17.2 | -, - | -, - |
| PA2508 | *catC* | 4.4 | 17.2 | -, - | -, - |
| PA2509 | *catB* | 3.9 | 13.1 | -, - | -, - |
| PA2587 | *pqsH* | 20.5 | 21.8 | 20.9, 18.5 | 5.2, 13.8 |
| PA2588 |  | 11.0 | 8.4 | 7.7, 5.3 | -, 4.9 |
| PA2589 |  | 3.7 | 3.2 | 4.5, - | -, - |
| **PA2591** | *vqsR* | 14.2 | 15.0 | 20.3, 16.8 | 8.8, 38.0 |
| PA2592 |  | 6.3 | 5.6 | 9.3, 7.2 | 3.0, 4.4 |
| PA2593 | *qteE* | 4.3 | 3.6 | 5.0, 4.8 | -, - |
| PA2664 | *fhp* | 11.9 | 11.0 | -, - | -, - |
| PA2939 |  | 10.8 | 21.1 | 26.4, 14.1 | 4.1, - |
| PA3104 | *xcpP* | 3.3 | 3.3 | -, 2.9 | -, - |
| **PA3326** | *azeA/clpP2* | 4.2 | 5.0 | 30.0, 15.7 | 10.6, 24.8 |
| **PA3327** | *azeB* | 5.9 | 10.1 | 34.6, 45.0 | 7.3, 36.4 |
| **PA3328** | *azeC* | 6.4 | 14.4 | 62.0, 47.9 | 7.7, 51.2 |
| **PA3329** | *azeD* | 5.6 | 14.3 | 25.0, 26.3 | 4.6, 20.1 |
| **PA3330** | *azeE* | 11.4 | 12.9 | 58.3, 50.8 | 7.0, 68.8 |
| PA3331 | *azeF* | 5.1 | 6.2 | 18.1, 16.2 | 3.0, 17.2 |
| **PA3332** | *azeG* | 6.3 | 22.1 | 53.6, 45.4 | 5.0, 80.0 |
| **PA3333** | *azeH/fabH2* | 3.6 | 8.3 | 42.1, 36.3 | 3.7, 12.3 |
| **PA3476** | *rhII* | 23.7 | 27.1 | 26.6, 28.9 | 19.3, 41.1 |
| PA3477 | *rhIR* | 7.5 | 6.8 | 8.4, 6.4 | -, 5.8 |
| **PA3479** | *rhIA* | 3.1 | 3.7 | 27.7, 18.8 | 7.8, 17.6 |
| PA3518 |  | 3.4 | 10.1 | -, - | -, - |
| PA3519 |  | 8.4 | 16.9 | -, - | -, - |
| PA3520 |  | 4.8 | 3.1 | 4.2, 3.3 | -, - |
| PA3535 | *eprS* | 22.0 | 20.0 | 8.7, 10.8 | -, 6.5 |
| **PA3724** | *lasB* | 53.5 | 63.8 | 51.6, 39.7 | 14.3, 23.2 |
| PA3904 | *PAAR4* | 22.3 | 44.8 | 18.4, 28.2 | 8.2, 28.1 |
| PA3905 | *tecT* | 27.0 | 29.7 | 31.1, 22.4 | -, 12.0 |
| PA3906 | *co-tecT* | 34.4 | 25.7 | 15.2, 15.4 | -, 5.2 |
| PA3907 | *tseT* | 23.2 | 30.7 | 20.3, 25.6 | -, 6.5 |
| PA3908 | *tsiT* | 24.2 | 18.9 | 24.1, 16.5 | -, 4.1 |
| PA3915 | *moaB1* | 4.3 | 4.3 | -, - | -, - |
| PA4117 | *bphP* | 4.8 | 3.8 | 4.5, 3.7 | -, 3.1 |
| PA4128 | *hcpH* | 4.2 | 6.9 | 24.5, 25.3 | 4.7, - |
| PA4129 |  | 5.8 | 9.0 | 46.6, 27.6 | 8.4, - |
| PA4130 | *nirA* | 6.0 | 7.8 | 37.0, 26.6 | 12.8, - |
| PA4131 |  | 5.4 | 4.8 | 27.7, 18.4 | 22.0, 46.0 |
| PA4132 |  | 4.1 | 3.7 | 16.1, 10.6 | 4.4, - |
| PA4133 |  | 10.2 | 16.7 | 38.3, 25.8 | 5.2, - |
| PA4134 |  | 2.9 | 5.1 | 11.3, 7.4 | -, - |
| PA4139 |  | 37.4 | 35.9 | 3.7, 6.3 | 4.0, - |
| PA4140 |  | 18.4 | 16.7 | -, - | 2.9, - |
| **PA4141** |  | 3.7 | 5.6 | 8.0, 6.9 | 4.1, 8.7 |
| PA4142 |  | 5.2 | 5.6 | 3.1, 3.5 | -, - |
| PA4143 |  | 3.5 | 3.3 | -, - | -, - |
| PA4175 | *piv* | 8.7 | 10.4 | 3.9, 3.2 | -, - |
| PA4190 | *pqsL* | 4.1 | 4.9 | 3.7, 4.4 | -, 3.1 |
| PA4589 |  | 3.2 | 2.9 | -, - | -, - |
| PA4590 | *pra* | 8.7 | 9.1 | 5.2, 4.1 | -, 4.6 |
| PA4637a |  | 3.7 | 6.7 | 3.9, 2.8 | -, - |
| PA4677 |  | 4.4 | 4.4 | 8.6, 7.7 | 3.4, 9.8 |
| PA4778 | *cueR* | 4.2 | 3.5 | 5.2, 4.5 | -, 3.5 |
| PA4869 |  | 4.1 | 3.8 | 2.8, - | -, - |
| PA5058 | *phaC2* | 3.6 | 3.6 | 4.6, 2.9 | -, - |
| PA5180 |  | 2.9 | 4.1 | -, - | -, - |
| PA5181 |  | 3.5 | 3.7 | -, - | -, - |

<sup>a</sup>Data are published in (Smalley *et al.*, 2022), RNAseq values are from Population (pop) D and E, respectively. The 5-day adenosine-casein populations were transferred daily for five days and the 160-day populations were transferred for 160 days prior to growth in buffered LB for RNAseq analysis. Dashes indicate fold change was <2.8.
<sup>b</sup>Locus tags are from https://www.pseudomonas.com (Winsor *et al.*, 2016).
<sup>c</sup>Red text shows the only genes that were not QS activated in the $\Delta pqsR$ mutant compared to the PAO1 WT. These include the first four genes in the *pqs* operon. The last gene, *pqsE* does not appear on the list because it did not show a >2.8 activation in cells without lactonase compared to cells grown with lactonase.
<sup>d</sup>Note the $\Delta pqsR$ mutant strain lacks ~95% of the *pqsR* open reading frame.
<sup>e</sup>Locus tags in bold are those designated as members of the RhIR-dependent core as defined elsewhere (Asfahl *et al.*, 2022).
<sup>f</sup>There are two copies of *phzA* and *phzB* on the *P. aeruginosa* PAO1 chromosome. The RNAseq analysis does not discriminate between the two and by default calls them PA1899 and PA1900, respectively. The results include transcripts of *phzA1* (PA4210) and *phzA2* (PA1899) combined, and *phzB1* (PA1900) and *phzB2* (PA4211) combined. These transcripts were not considered in Smalley *et al.* (Smalley *et al.*, 2022).

The lists of QS-activated genes in the WT and the Δ*pqsR* mutant were strikingly similar (Table 1) and differed only the differential expression of *pqsR* (which was deleted in the mutant) and the *pqsA-E* operon. This result provides evidence that the reductive evolution of the QS regulon in the populations evolved on casein plus adenosine is not the result of a simple mutation that inactivates *pqsR*. But if *pqsR* is required to activate expression of *pqsE*, and PqsE is required for full expression of at least several genes activated by RhlR, then why we did not observe the loss of genes other than those in the *pqsA-E* operon in the QS transcriptome comparisons of WT and the Δ*pqsR* cells, or at least a diminution of the levels of QS activation? There are several possible explanations. For example, we know that the RhlR regulon is activated later than the LasR regulon, at least in part, because *rhlR-rhlI* are activated by LasR in PAO1 (Gilbert *et al*., 2009, Latifi *et al*., 1996). We note that a recent report identified 20 genes in the “core” RhlR regulon, genes activated by RhlR in several *P. aeruginosa* clinical isolates (Asfahl *et al*., 2022). Of this core 19 were QS-activated as assessed by our comparative transcriptomics approach (locus tags in bold text in Table 1). It may also be that *pqsE* is sufficiently expressed in the *pqsR* mutant under the conditions of our transcriptomics experiment, or it could be that, under the specific conditions of our experiments, RhlR is sufficiently active in the absence of PqsE. These possibilities are not mutually exclusive.

### QS gene activation in a Δ*pqsE* mutant

To begin to understand why the QS regulons of WT and a PqsR mutant were so similar, we first wanted to address the question of whether *pqsE* might be expressed at sufficient levels to chaperone RhlR even in a PqsR mutant. To address this issue, we constructed a *pqsE* deletion mutant and compared the QS regulon of the mutant to that of WT. Of the 129 QS-activated genes in the WT, 124 were also activated in the Δ*pqsE* mutant. The five genes lost from the QS regulon in the PqsE mutant are listed in Table 2. From this analysis we conclude that, under the conditions of our experiment, the minimal influence of PqsR on the QS regulon does not result from PqsR-independent expression of *pqsE*. Our results do not address the question of whether the *pqsR* deletion mutant expresses *pqsE* sufficiently to serve as a RhlR chaperone. In fact, all five PqsE-dependent transcripts in Table 2 showed similar levels of QS activation in WT and the *pqsR* deletion mutant (see Table 1).

**Table 2.**
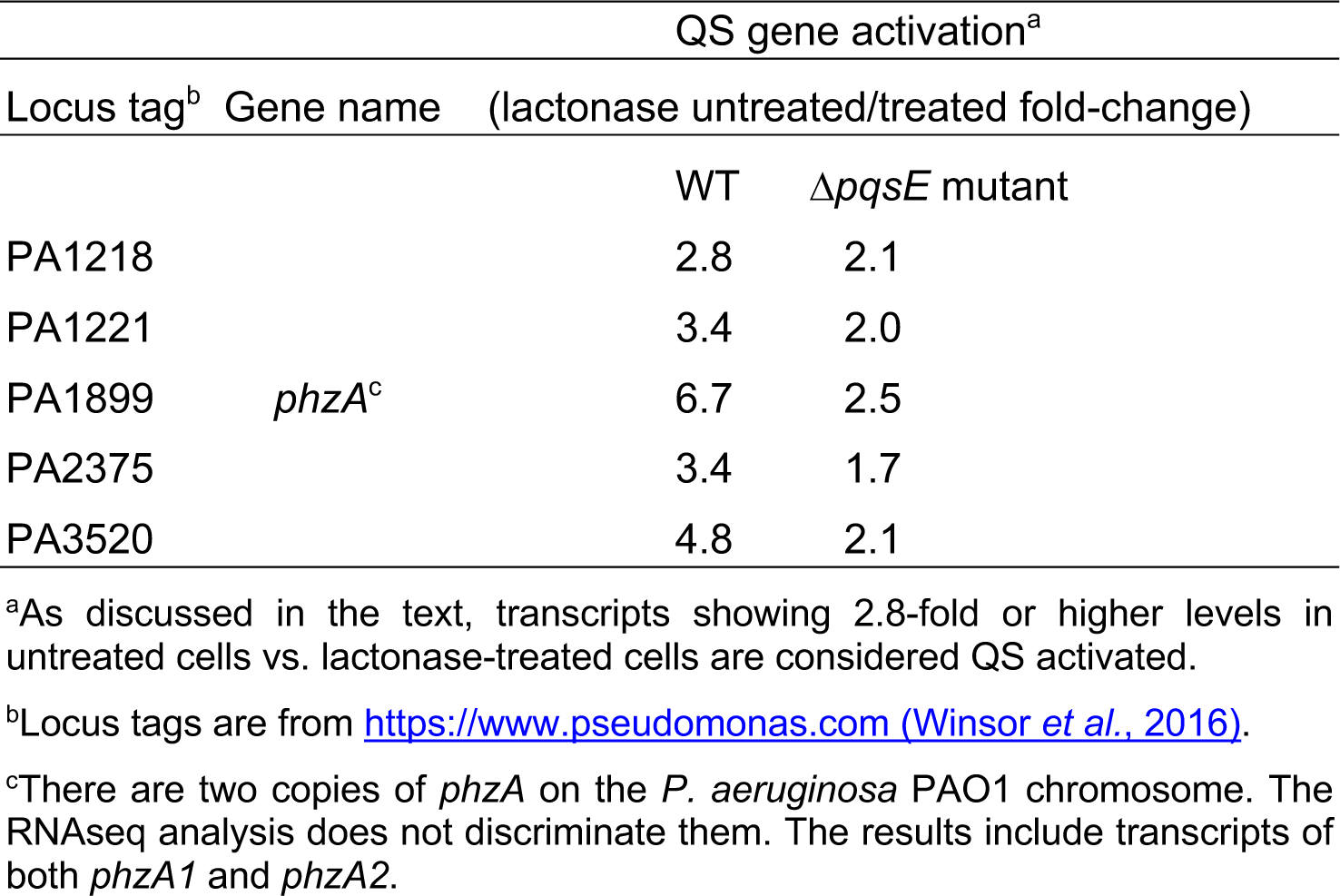
Transcripts activated by QS in WT but not in the Δ*pqsE* mutant.

| Locus tag<sup>b</sup> | Gene name | QS gene activation<sup>a</sup> |  |
| --- | --- | --- | --- |
|  |  | (lactonase untreated/treated fold-change) |  |
| | | WT | $\Delta pqsE$ mutant |
| PA1218 |  | 2.8 | 2.1 |
| PA1221 |  | 3.4 | 2.0 |
| PA1899 | *phzA*<sup>c</sup> | 6.7 | 2.5 |
| PA2375 |  | 3.4 | 1.7 |
| PA3520 |  | 4.8 | 2.1 |

<sup>a</sup>As discussed in the text, transcripts showing 2.8-fold or higher levels in untreated cells vs. lactonase-treated cells are considered QS activated.
<sup>b</sup>Locus tags are from https://www.pseudomonas.com (Winsor *et al.*, 2016).
<sup>c</sup>There are two copies of *phzA* on the *P. aeruginosa* PAO1 chromosome. The RNAseq analysis does not discriminate them. The results include transcripts of both *phzA1* and *phzA2*.

One of the few transcripts affected by PqsE was *phzA* (Table 2). We examined *phzA* expression further for two reasons. First, there are two copies of the *phzA* gene in *P. aeruginosa*, *phzA1* (PA4210) and *phzA2* (PA1899). The *phzA1* operon is QS-dependent (Whiteley & Greenberg, 2001). The *phz* genes are responsible for synthesis of the blue pigment pyocyanin (Mavrodi *et al*., 2001), and production of this pigment is QS-(Wagner *et al*., 2003, Schuster *et al*., 2003) and PqsE-dependent (Groleau *et al*., 2020, Keegan *et al*., 2023). To examine the influence of PqsE on the *phzA1* operon specifically, we performed RT-PCR with *phzA1*-specific primers and we also used a reporter with the *phzA1* promoter fused to a gene encoding the Green Fluorescent Protein (GFP) (Fig. 1). Results of the two experiments were consistent in that they confirmed that PqsE was required for QS-dependent activation of the *pqsA1* operon.

**Figure 1.**
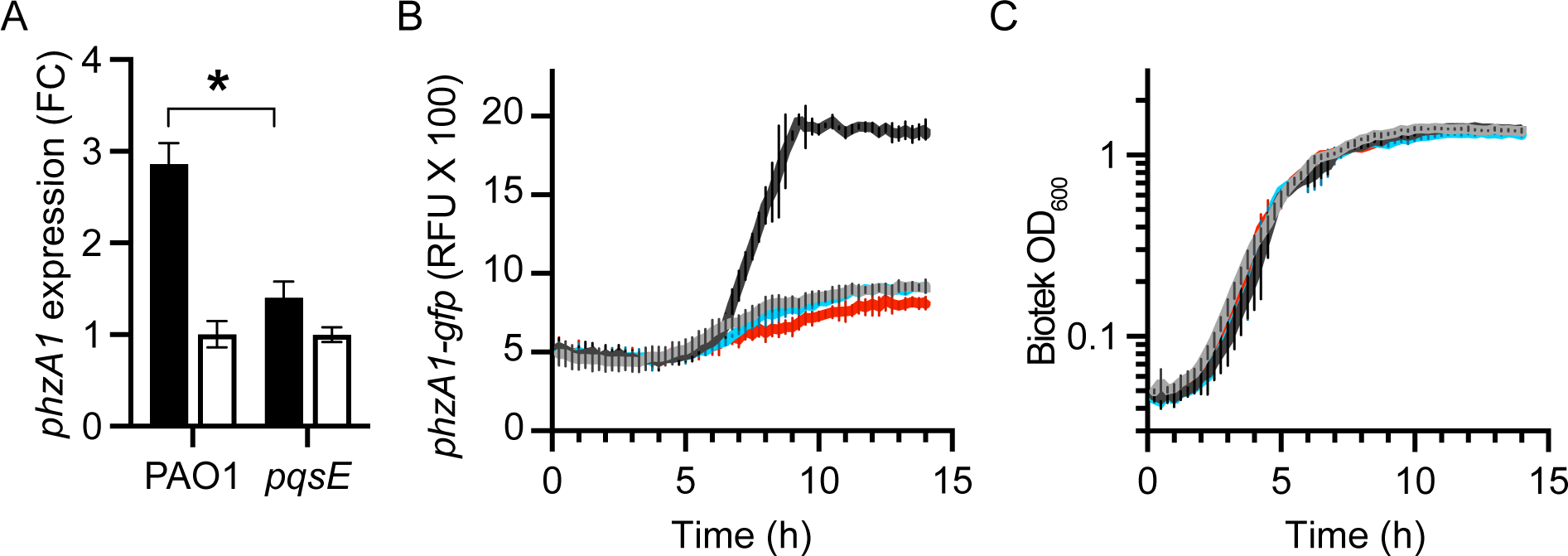
Expression of *phzA1* in WT and a *pqsE* deletion mutant. (A) Relative levels of *phzA1* transcript as measured by qRT-PCR reported as fold-change (FC) relative to the housekeeping gene *rplU* as described in METHODS. RNA was isolated from cells harvested when the culture densities reached an OD_600_ of 2.0. Cells were grown in buffered LB with (white) or without (black) AiiA lactonase as indicated. Results are shown as means of two biological replicates (each with two technical replicates) with ranges, the asterisk indicates a significant difference in an unpaired t-test (*p*<0.05). (B) GFP fluorescence and (C) growth (OD_600_) as measured by Biotek plate reader in PAO1 WT (black), a *pqsE* deletion mutant (blue), and a *rhlR* deletion mutant (red) all harboring a *phzA1-gfp* reporter; PAO1 harboring the vector control (grey) is also included. For these experiments growth was in 0.3 ml volumes of buffered LB in wells of a 48-well microtiter dish. Graphed in B and C are the means of three biological replicates and the error bars represent the standard error of the mean.

We were interested in the four additional genes, which showed reduced QS activation in our *pqsE* deletion mutant (Table 2). The role of PqsE as a RhlR chaperone is complicated in that RhlR activation of a few genes shows strong dependence on PqsE, whereas RhlR shows a modest PqsE-independent ability to activate other genes, and RhlR activation of yet other genes seems minimally influenced by PqsE (Keegan *et al*., 2023). Do the other genes listed in Table 2 show strong dependence on PqsE? Among the four genes, PA1218 and PA1221 are likely part of a single transcriptional unit transcribed from the PA1221 promoter (Wurtzel *et al*., 2012) and predicted by anti-SMASH to be non-ribosomal peptide synthetase-like/betalactone genes (Blin *et al*., 2023). PA2375 is monocistronic and codes for a small (131 amino acid) polypeptide of unknown function. This gene is embedded in a cluster of Type VI secretion genes and may code for an antitoxin. PA3520 appears to be monocistronic and its product has been annotated as a periplasmic binding protein. We constructed reporter plasmids with the promoters of PA1221, 2375 and 3520 fused to a gene coding for GFP and measured GFP fluorescence during growth of WT, the *pqsE* deletion mutant, and a *rhlR* deletion mutant during growth in microtiter plate dishes. Our results show that, like *phzA1* expression, expression from the PA1221 and PA3520 promoters was severely constrained in the PqsE mutant, whereas expression from the PA2375 promoter was similar in WT, the *pqsE* deletion mutant, and the *rhlR* deletion mutant. (Fig. 2). We conclude that under the conditions of these experiments PA1221 and PA3520 require both RhlR and PqsE for activation, and PA2375 does not appear to be RhlR- or PqsE-dependent.

**Figure 2.**
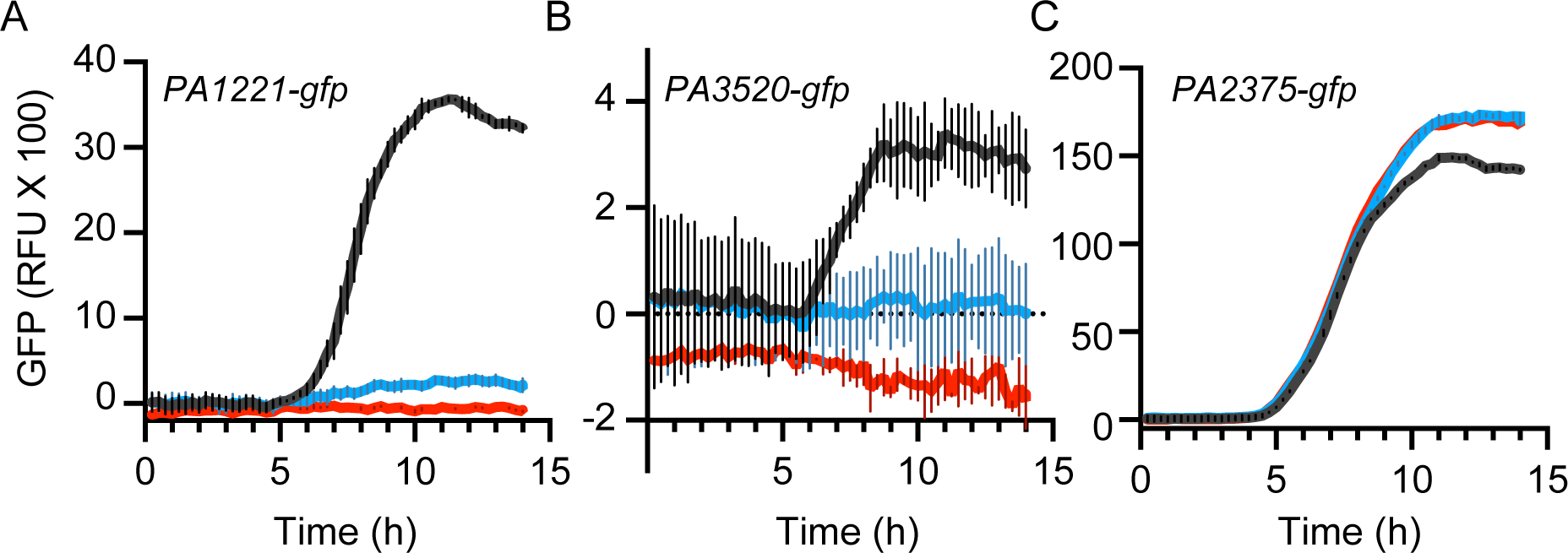
Expression from the PA1221 and PA3520 promoters, but not the PA2375 promoter, are RhlR- and PqsE-dependent. Fluorescence in the PAO1 WT (black), *pqsE* deletion mutant (blue), or *rhlR* deletion mutant (red) harboring a *PA1221-gfp* fusion (A), *PA3520-gfp* fusion (B), or *PA2375-gfp* fusion (C). Fluorescence of cells expressing the plasmid vector was subtracted. Cells were grown in 0.3 ml volumes of buffered LB in wells of a 48-well microtiter dish. Results are plotted as the mean of three or four biological replicates and the error bars represent the standard error of the mean.

The transcriptomics results shown in Table 1 suggest that RhlR-activated genes, including those expressed from the PA1221 promoter and the PA3520 promoter, were not suppressed in the PqsR mutant and, apart from those expressed from the promoters of *phzA1*, PA1221 and PA3520, transcriptomics did not support the idea that PqsE was important for their expression (Table 2). We reasoned that this apparent conflict with reports that PqsE binding to RhlR is important for its activity might be related to specific experimental details. Perhaps the conflict results from the use of different *P. aeruginosa* strains or from the use of different culture conditions. In an effort to resolve the inconsistency, we constructed reporter plasmids with two RhlR-dependent genes described elsewhere as showing different levels of PqsE dependence (Keegan *et al*., 2023), *rhlA* fused to *gfp* and *hcnA* fused to *gfp*, and measured GFP fluorescence of cells containing these reporters during growth in microtiter dish experiments (Fig. 3). There was clear PqsE-independent induction of GFP, as expression in WT was about four times that of the PqsE mutant.

**Figure 3.**
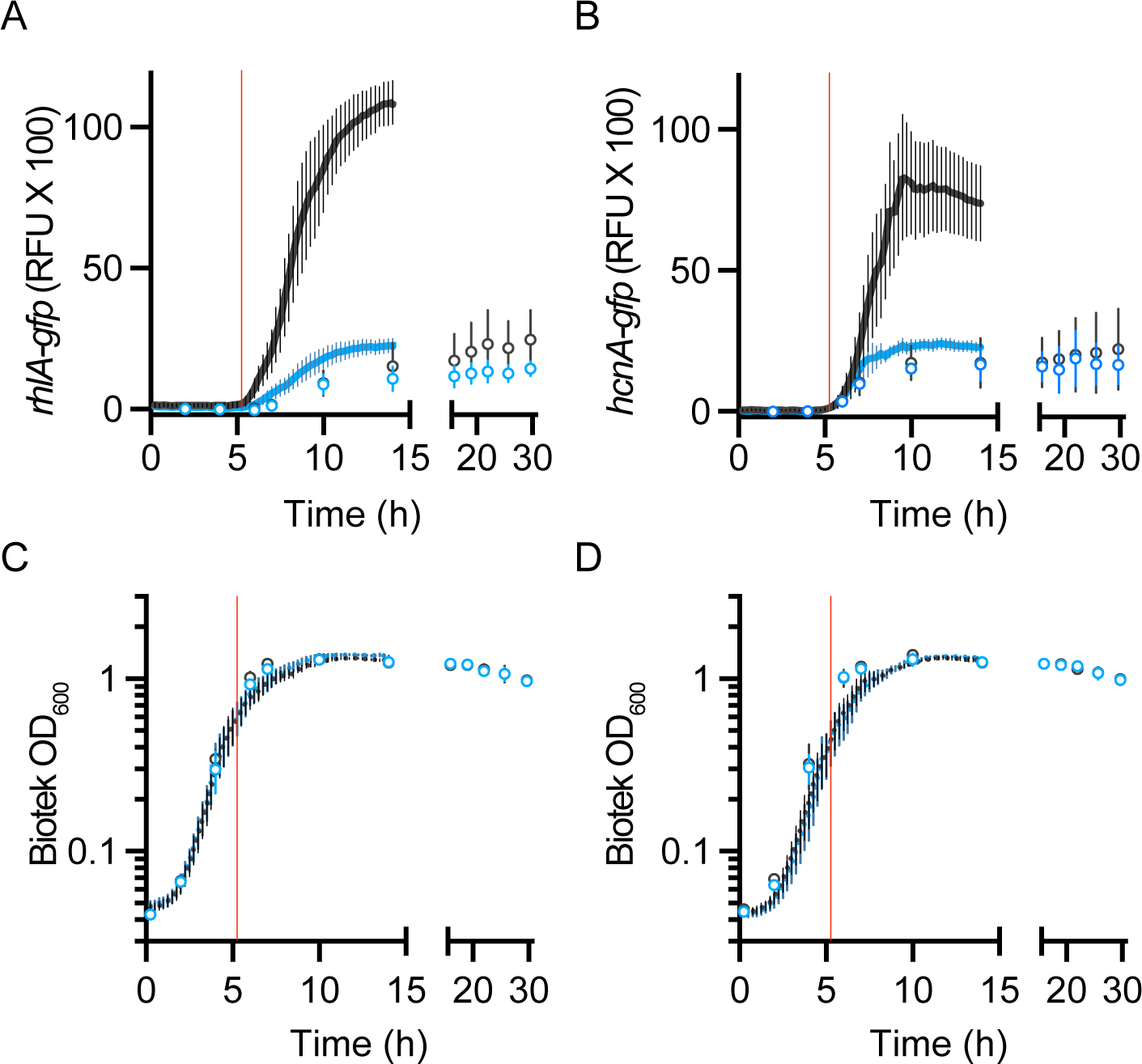
Expression of *rhlA-gfp* (A) or *hcnA-gfp* (B) reporters in the WT (black) and *pqsE* deletion mutant (blue) grown in shaking flasks (open circles) or microtiter dish wells (solid lines). Fluorescence of cells with the plasmid vector was subtracted. Growth of each culture condition (well-aerated flask and 48-well plate) for the *rhlA-gfp* (C) and *hcnA-gfp* (D) containing strains was measured by optical density at 600 nm in the Biotek plate reader (Biotek OD_600_). Results are plotted as the mean of two biological replicates (error bars represent the range) in well-aerated flasks or three to four biological replicates (error bars represent the standard error of the mean in 48-well plates. The red line estimates the time when the Genesys OD_600_ reading would be 2.0 (equivalent to an OD_600_ in the microtiter dishes of about 0.6), which is when we harvested RNA for transcriptomics (Tables 1 and 2).

In our microtiter dish experiments, expression of the *rhlA-gfp* reporter and a *hcnA-gfp* reporter depend on PqsE (Fig. 3), whereas there is no apparent influence of PqsE on expression of *rhlA* or *hcn* genes as assessed by RNAseq in our well-aerated flask-grown cells (Table 1). To assess whether the conflicting results were due to the use of RNAseq vs. reporter technology or to the different growth conditions (microtiter dish vs. flask growth) we performed reporter experiments with cells containing the *rhlA-gfp* reporter plasmid grown in flasks (Fig. 3A, open circles). Whereas differences in GFP levels can be seen in microtiter dish cultures by eight hours of growth, in late logarithmic phase, in the shaking flask cultures any differences in GFP levels were small throughout logarithmic phase and were evident only in late stationary phase, after about twenty hours of growth. Notably, at a Genesys OD_600_ reading of 2 (red lines in Fig. 3), the density of cultures used for RNAseq experiments, the differences in GFP fluorescence of the WT and the *pqsE* deletion mutant were negligible. We note that, after cessation of logarithmic growth, GFP fluorescence continues to rise in both the WT and the *pqsE* deletion mutant, but more so, about twice as much, in the WT. These results help resolve inconsistencies in literature in that experimental design and conditions can reveal different aspects of QS control of gene expression.

## DISCUSSION

We tested the hypothesis that the reductive evolution of genes controlled by QS in *P. aeruginosa* observed after long term QS-dependent growth was a result of a mutation that inactivates *pqsR* (Smalley *et al*., 2022). Results from our transcriptome analysis show that the reported reduction in the QS regulon of *P. aeruginosa* PAO1 cannot be explained by a simple *pqsR* mutation (Table 1). The reduction in the QS regulon must be the consequence of other factors.

Our findings led us to probe the relationship between *pqsR* and the PqsR-activated *pqsE* gene to QS in strain PAO1. Studies with strain PA14 show that PqsE physically interacts with RhlR (Simanek *et al*., 2022, Feathers *et al*., 2022) and influences RhlR binding to target promoters, and RhlR activation of those promoters (Keegan *et al*., 2023). The influence of PqsE on RhlR binding and gene activation varies among different RhlR-dependent promoters. In PA14 PqsE has very little influence on some RhlR-dependent promoters and RhlR activation of other promoters relies heavily on PqsE. Dependence of PqsE for RhlR at other promoters seems to fall on a continuum from very strong to very weak (Keegan *et al*., 2023). We found that QS-dependent gene expression in WT PAO1 and a *pqsR* deletion mutation were similar with the main difference being that the PqsR mutant did not exhibit QS activated expression of the *pqsA-E* operon (Table 1). The lack of *pqsA-E* induction was expected because this operon is indirectly activated by LasR by virtue of LasR activation of *pqsR* expression (Gallagher *et al*., 2002). If, however, *pqsE* expression is dependent on PqsR, and PqsE is needed for activation of many RhlR-dependent genes, why were these genes not captured in our RNAseq analysis? Of note, the QS-dependent *phzA1* operon shows nearly absolute dependence on PqsE. We did not capture any difference in *phzA* expression in WT vs. the PqsR mutant. There are two *phzA* operons in *P. aeruginosa* and our transcriptomics analysis does not discriminate between them. Thus, from the transcriptomics analysis in Table 1 we cannot rule out the possibility that *phzA1* transcription was influenced by the *pqsR* mutation. Our follow up experiments revealed that RhlR-dependent transcription from the *phzA1* promoter is PqsE dependent (Table 2). In fact, we identified four PqsE-dependent transcriptional units by comparing transcriptomes of WT and a PqsE deletion mutant (Table 2), three of which, including the *pqsA1* operon, were confirmed as PqsE-dependent by RT-PCR and *gfp* fusion experiments (Fig. 1 and 2).

The transcriptomics results appear in conflict with previous reports about the involvement of PqsE in RhlR-dependent gene activation (Simanek *et al*., 2022). We sought to resolve the conflict by constructing reporter plasmids with *gfp* fused to the RhlR-dependent *rhlA* and *hcnA* promoters (Fig. 3). By following GFP fluorescence during growth in either microtiter dish wells or in shaking flask cultures we learned that PqsE is important for RhlR activation of these promoters in late logarithmic phase in microtiter dish wells, but its role in RhlR activation of these promoters in shaking flask cells in minimal in late logarithmic and early stationary phases, and manifests itself much later in stationary phase, long after growth has ceased. Notably, in flask cultures at an OD of 2, the OD at which cells are harvested for RNAseq studies (as measured using a Genesys spectrophotometer), PqsE has little influence on RhlR activation of the *rhlA* and *hcnA* promoters. Our experiments help clarify inconsistencies regarding the role of PqsE in RhlR-dependent gene activation. For genes such as *rhlA* and *hcnA*, where expression is not absolutely abolished by a *pqsE* deletion, we find PqsE provides only a 3-5-fold stimulation of transcription (Fig. 3). We do not yet understand why PqsE enhancement of RhlR gene activation for at least some genes depends on environmental conditions. Differences between microtiter dish growth and shaking flask growth might include oxygen availability and cellular redox state, cells in microtiter dish wells might aggregate where this might be discouraged by growth in flasks with vigorous shaking. Finally, there is evidence that cellular levels of RhlR and concentrations of C4-HSL, the QS signal for RhlR, can influence the level of PqsE stimulation of RhlR activity (Keegan *et al*., 2023). It is not unlikely that levels of these factors might be influenced by growth conditions.

## MATERIALS and METHODS

### Bacterial strains and culture conditions

Bacterial strains and plasmids are described in Table 3. *Escherichia coli* was grown in Lennox Broth [LB,(Lennox, 1955)], and *P. aeruginosa* was grown in LB buffered with 50 mM 3-(N-morpholino) propane sulfonic acid, (MOPS, pH 7.0) (buffered LB). For all the experiments with *P. aeruginosa* we used previously described growth conditions (Smalley *et al*., 2022). Briefly, precultures were inoculated from an overnight culture to an initial optical density at 600 nm (OD_600_, measured using a Genesys spectrophotometer) of 0.01 in 5 mL of buffered LB in 18 mm tubes. When precultures reached logarithmic phase (OD_600_ 0.1-0.4), they were used as inocula at a starting OD_600_ of 0.005 in buffered LB at volumes of either 25 mL of in 250 mL baffled flasks or 300 μl in wells of 48-well microtiter plates. Where indicated, purified AiiA lactonase (100 μg per mL), or an equal volume of buffer (20 mM Tris-HCL, 100 mM NaCl, pH 7.4, 10% glycerol), was added as described previously (Smalley *et al*., 2022). Growth was at 37°C. Flasks were shaken continuously on a rotary shaker at 225 rpm and microtiter plates were shaken continuously with a double orbital setting. When strains carried plasmids, gentamicin (10 μg/ml) was included.

**Table 3.** Strains and plasmids used in this study.

| Strain or plasmid | Relevant characteristic | Reference or source |
| --- | --- | --- |
| **Strains** |  |  |
| *E. coli* |  |  |
| DH5α | *fhuA2 Δ(argF-lacZ)U169 phoA glnV44 Φ80 Δ(lacZ)M15 gyrA96 recA1 relA1 endA1 thi-1 hsdR17* | Invitrogen |
| *P. aeruginosa* |  |  |
| MPAO1 | Wild-type strain | (Jacobs *et al.*, 2003) |
| Δ*pqsR* | PAO1 *pqsR*<sub>Δ54-958</sub> deletion mutant | This study |
| Δ*pqsE* | PAO1 *pqsE*<sub>Δ10-903</sub> deletion mutant | This study |
| **Plasmids** |  |  |
| pBBR-*gfp* | Promoterless *gfp* transcriptional reporter; Gm<sup>R</sup>. | (Smalley *et al.*, 2022) |
| pP<sub>phzA1</sub>-*gfp* | pBBR-*gfp* with -395 to +79 bp of the *phzA1* promoter; Gm<sup>R</sup> | This study |
| pP<sub>PA1221</sub>-*gfp* | pBBR-*gfp* with -475 to +60 bp of the PA1221 promoter; Gm<sup>R</sup> | This study |
| pP<sub>PA3520</sub>-*gfp* | pBBR-*gfp* with -393 to +65 bp of the PA3520 promoter; Gm<sup>R</sup> | This study |
| pP<sub>PA2375</sub>-*gfp* | pBBR-*gfp* with -293 to +81 bp of the PA2375 promoter; Gm<sup>R</sup> | This study |
| pP<sub>rhIA</sub>-*gfp* | pBBR-*gfp* with -500 to +31 bp of the *rhIA* promoter; Gm<sup>R</sup> | (Kostylev *et al.*, 2023) |
| pP<sub>hcnA</sub>-*gfp* | pBBR-*gfp* with -256 to +52 bp of the *hcnA* promoter; Gm<sup>R</sup> | This study |
| pEXG2 | Allelic exchange vector with pBR origin; *sacB*, Gm<sup>R</sup> | (Rietsch *et al.*, 2005) |
| pEXG2-*pqsR*-KO | pEXG2 with flanking DNA sequence (448 bp upstream, 415 bp downstream) used to create the *pqsR*<sub>Δ54-958</sub> deletion allelic exchange vector; *sacB*, Gm<sup>R</sup> | This study |
| pEXG2-*pqsE*-KO | pEXG2 with flanking DNA sequence (501 bp upstream, 499 bp downstream) used to create the *pqsE*<sub>Δ10-903</sub> deletion allelic exchange vector; *sacB*, Gm<sup>R</sup> | This study |

**Abbreviations:** Gm<sup>R</sup>, gentamicin resistance

### Mutant and plasmid construction

To create the Δ*pqsR* (deleted nucleotides 54-968 of the 999-bp open reading frame) and Δ*pqsE* (deleted nucleotides 10-903 of the 906-bp open reading frame) mutants, we used a homologous recombination-based two-step allelic exchange with sucrose counterselection approach as described previously (Kostylev *et al*., 2019, Hmelo *et al*., 2015). We cloned about 450 bp of flanking DNA into the suicide vector pEXG-2 (Rietsch *et al*., 2005) to create the appropriate knock-out constructs (Table 3), which used to electrotransform *P. aeruginosa* cells (Choi & Schweizer, 2005). Transformants were plated on agar plates with gentamicin (100 μg/ml) for the initial selection. The PA1221, PA2375 and PA3520 reporter plasmids (pP*_PA1221-gfp_*, pP*_PA2375-gfp_* and pP*_PA3520-gfp_*) were constructed by cloning the promoter regions (detailed in Table 3) into the plasmid pBBR-*gfp* (Smalley *et al*., 2022) using an assembly protocol described previously (Kostylev et al., 2015). Plasmid constructs were confirmed by whole plasmid sequencing (Plasmidsaurus, Eugene, OR).

### RNAseq

For RNA extraction, cells were grown as 25-ml cultures in 125-ml shaking flasks as described above and previously (Smalley *et al*., 2022). Once cultures reached an OD_600_ of 2 [as determined by a Genesys spectrophometer (ThermoFisher, Waltham, MA)], a 2-ml volume of cells were pelleted by centrifugation, snap frozen using liquid nitrogen, and the frozen pellets were stored at -80°C. RNA samples were prepared by using RNeasy Mini kits (Qiagen), followed by DNA depletion with Turbo DNase (Ambion) as described previously (Smalley *et al*., 2022). The DNA-free RNA was concentrated by MinElute Cleanup kit (Qiagen). The rRNA depletion, library preparation and sequencing were performed by Azenta formerly Genewiz. Library samples runs were performed with paired-end 2 x 150 bp read lengths. Trim Galore! (Babraham Bioinformatics, Cambridge UK) was used to trim adapters prior to alignment against the PAO1 reference genome (accession NC_002516) using StrandNGS version 3.3.1 (Strand Life Sciences, Bangalore, India). DESeq2 (Love et al., 2014) was used for differential expression analysis, using the Benjamini-Hochberg adjustment for multiple comparisons and a false-discovery rate a = 0.05. QS-activated regulons were called by imposing a 2.8-fold minimal fold change threshold (this is 1.5-fold on a log2 scale). Raw sequencing reads and count matrices for RNAseq experiments summarized in Tables 1 and 2 were deposited into the NCBI Sequence Read Archive as Bioproject PRJNA1063357 (Biosamples SAMN40421732 thru SAMN40421743) and GEO accession GSE261453.

### qRT-PCR

RNA extraction and DNA depletion was performed using the protocol described above. We used an iScript Select cDNA Synthesis Kit (BioRad, Hercules, CA) to generate cDNA. For qPCR we used SsoAdvanced SYBR Green Supermix (BioRad, Hercules, CA). The *phzA1*-specific product (forward primer GGCTATTGCGAGAACCACTA, reverse primer CAATGCACGCAGTTTCTGTA) was normalized to the housekeeping gene *rplU* product (forward primer CAAAGTCACCGAAGGCGAAT, reverse primer ACGCTTCATGTGGTGCTTAC) as described previously (Cruz *et al*., 2020). In each case, the fold-change in gene expression is relative to its expression in the same strain but with lactonase addition (QS active vs. QS inactive). This was calculated using the 2^-1ΔΔCt^ method (Livak & Schmittgen, 2001).

### Reporter assays

For reporter experiments growth was in microwell plates [48 well, clear flat bottom plates item FB012930 (Fisher Scientific, Waltham MA)], containing 300 mL of culture in each well, incubated at 37°C with continuous readings in a Biotek Synergy H1 microplate reader (Winooski, VT), or in flask cultures as indicated. Optical density at 600 nm (OD_600_) and GFP fluorescence (489 nm excitation, 520 nm emission, gain 80) were measured in the microtiter plate reader (0.3 ml culture volumes in a 48-well plate) unless otherwise indicated.

## ACKNOWLEDGEMENTS

We thank Maureen Thomason, Indraneel Salukhe and Crystal Perez for sharing plasmids. This work was supported by NIH grant R35 GM136218 (to E.P.G.).

## Notes

### Competing Interest Statement

The authors have declared no competing interest.

